# Effects of pathogenic CNVs on physical traits in participants of the UK Biobank

**DOI:** 10.1101/355297

**Authors:** David Owen, Mathew Bracher-Smith, Kimberley Kendall, Elliott Rees, Mark Einon, Valentina Escott-Price, Michael J Owen, Michael C O’Donovan, George Kirov

## Abstract

**Background:** Copy number variants (CNVs) have been shown to increase risk for physical anomalies, developmental, psychiatric and medical disorders. Some of them have been associated with changes in weight, height, and other physical traits. As most studies have been performed on children and young people, these effects of CNVs in adulthood are not well established.

**Methods:** The UK Biobank recruited half a million adults who provided a variety of physical measurements. We called all CNVs from the Affymetrix microarrays and selected a set of 54 CNVs implicated as pathogenic (including their reciprocal deletions/duplications) and that were present in five or more persons. Linear regression analysis was used to establish their association with 16 physical traits, relevant to human health.

**Results:** 396,725 participants of white British or Irish descent (excluding first-degree relatives) passed our quality control filters. There were 214 CNV/trait associations significant at a false discovery rate of 0.1, most of them novel. These traits are associated with adverse health outcomes: e.g. increased weight, waist-to-hip ratio, pulse rate and body fat composition. Deletions at 16p11.2, 16p12.1, *NRXN1* and duplications at 16p13.11 and 22q11.2 produced the highest numbers of significant associations. CNVs at 1q21.1, 2q13, 16p11.2, 16p11.2 distal, 16p12.1, 17p12 and 17q12 demonstrated one or more mirror image effects of deletions versus duplications.

**Conclusions:** Carriers of many CNVs should be monitored for physical traits that increase morbidity and mortality. Genes within these CNVs can give insights into biological processes and therapeutic interventions.

## INTRODUCTION

Human height, weight and other anthropometric traits are highly heritable. Genetic factors contribute up to 80% of height (1,2), and genome-wide association studies (GWAS) suggest that the total additive effect of single nucleotide variants (SNVs) explain 56% of variance in height and 27% of body mass index (BMI) variability (3). Hand grip strength’s heritability has been estimated at 56% (4). Resting heart rate, systolic and diastolic blood pressure are also highly heritable, with estimates of 61%, 54% and 49% respectively (5).

While SNVs tend to have small effect sizes, large copy number variants (CNVs) have been shown to have profound effects on weight and height, with 16p11.2 deletions and duplications providing a striking example (6). Other recognised CNVs with large effect sizes on physical traits are deletions at distal 16p11.2, associated with obesity (7) and at 1q21.1, associated with microcephaly and short stature (8). Failure to thrive has been described in some carriers of 3q29 duplications (9) while short stature has been associated with the classical and distal 22q11.2 deletions (10). Individuals with severe early-onset obesity have an increased rate of large and rare deletions (11). Obesity can be a feature in rare syndromic monogenic disorders (12), which include several of the CNVs analysed in the current study: 1q21.1 deletion, 15q11-q13 duplication, 16p11.2 classic and distal deletions and 22q11.2 duplication.

Most of the published research has been based on children or young people referred for genetic testing for developmental delay, congenital malformations and autism spectrum disorders, i.e. some of the most affected individuals. Such individuals may not be typical of carriers of CNVs, and importantly for long term health outcomes, the impact of these CNVs in adults, especially in those who have not been diagnosed as CNV carriers in early life, is not well described. In addition, most reports focus on a single, or at most a limited number of CNVs, making it difficult to perform comparative studies of the impact of individual CNVs on physical measures. Another potential problem is that small sample sizes will miss more subtle changes in physical traits.

The largest study on CNVs and anthropometric measures was performed on 191,161 unrelated European adults (13). It assessed systematically the effect of CNVs on BMI, weight, height, and waist/hip ratio and was based on 25 component studies of the Genetic Investigation of Anthropometric Traits (GIANT) Consortium, combined with the first release of UK Biobank data, approximately a third of the total sample. A genome-wide analysis implicated seven CNV regions: 1q21.1 (distal part: 145-145.9Mb), 3q29 (two sub-regions), 7q11.23, chr11: 26.97-27.19Mb; 16p11.2 distal, 16p11.2 classic; chr18: 55.81-56.05Mb and 22q11.21 (Mb intervals are in hg18). These loci were associated at genome-wide significance with at least one of the four traits in a mirror effect model in which deletions and duplications affect the trait in opposite directions. No common CNVs were significantly associated with the traits, despite the higher statistical power to detect such associations.

Here we report an analysis of the full UK Biobank cohort. We tested 54 CNVs that have been proposed to be pathogenic and were carried by at least five participants. We analysed these CNVs for association with an extended set of 16 physical traits (Methods), that include anthropometric traits (weight, height, BMI, waist, hip, waist/hip ratio), and other physical traits relevant to human health: pulse rate, blood pressure, arm strength, peak expiratory volume, heel bone mineral density, and the fat percentage of legs, arms and trunk. These traits have been associated with adverse health outcomes and increased mortality (14–19).

## RESULTS

The 54 CNVs and 16 traits produce 864 phenotype/CNV associations (Supplementary Table 1). Of those, 272 were nominally significant (at p < 0.05), a much higher number than the 43 expected by chance. Using Benjamini-Hochberg’s false discovery rate (FDR), 214 of those were significant at 0.1 (marked in bold in Supplementary Table 1). Images of the changes in the physical traits associated with each CNV, and their 95% confidence intervals (95%CI), are shown in Supplementary Figure 1 and on our institutional website (http://kirov.psycm.cf.ac.uk/Physicalmeasurements:searchbyCNV).

Table 1 summarises the significant findings. It is restricted to CNVs that have at least one significant association at FDR = 0.1, with the direction of the effect indicated with + or −. To simplify the presentation, we grouped together the three fat % measures to indicate any change on arm, leg or trunk fat % measures and we don’t show waist and hip circumferences, as the waist/hip ratio contains this information. Figure 1 shows the distribution of changes for the three CNVs with the largest number of significant associations (13 each). The significance also depends also on sample size, so these are not necessarily the most pathogenic CNVs.

**Figure 1.**
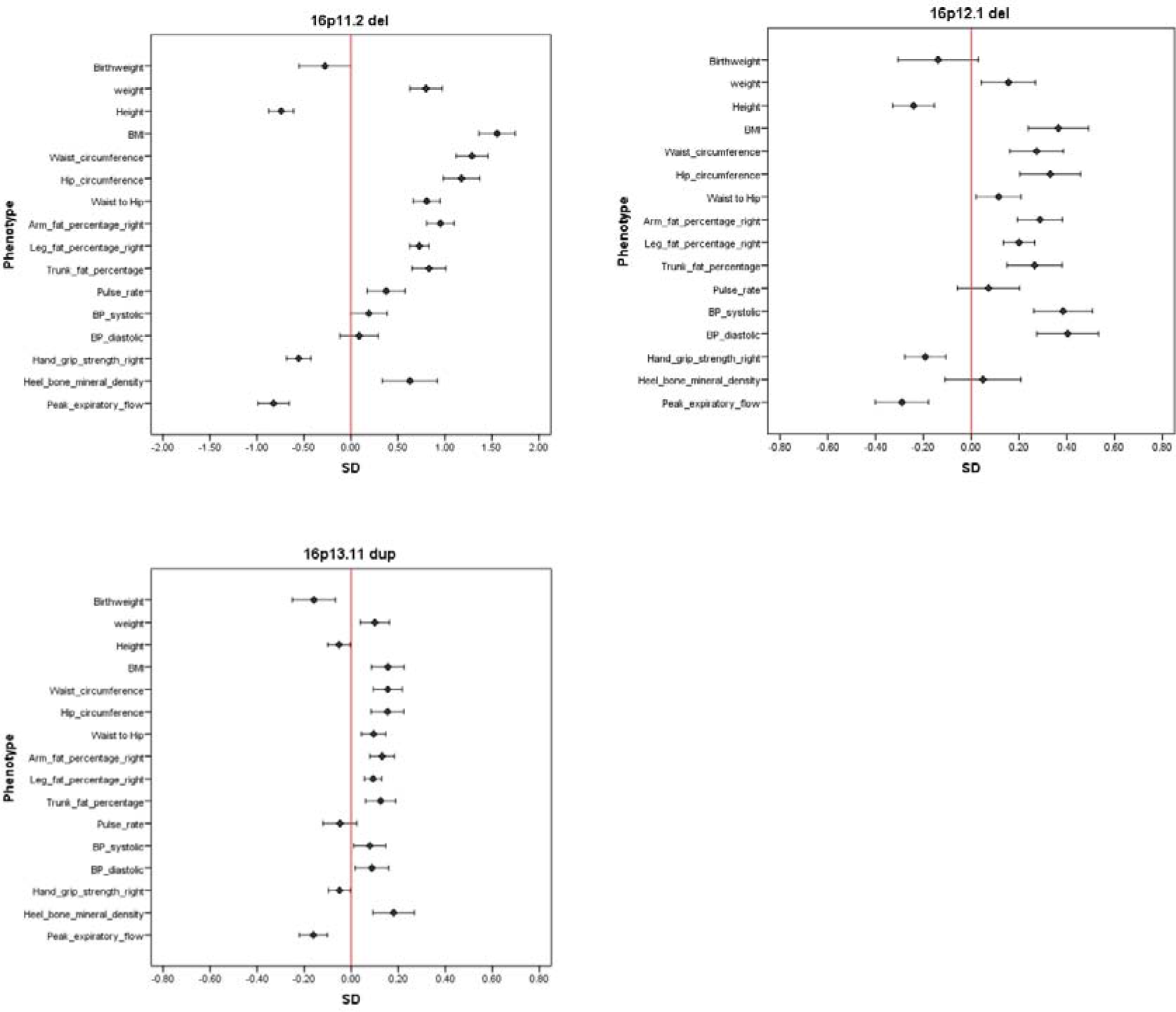
Effect sizes for the physical measurements for the three CNVs with the highest number of significant associations. Effects sizes means are the normalised (z-scored) values (standard deviation changes) after correction in linear regression analysis, with co-variates, produced with the *glm* function of R. The figures show the mean change in standard deviations (SD) and the 95% confidence intervals of the changes. a) 16p11.2 classic deletion,b) 16p12.1 deletion, c) 16p13.11 duplication. 95% CI are also shown.

**Table 1.**
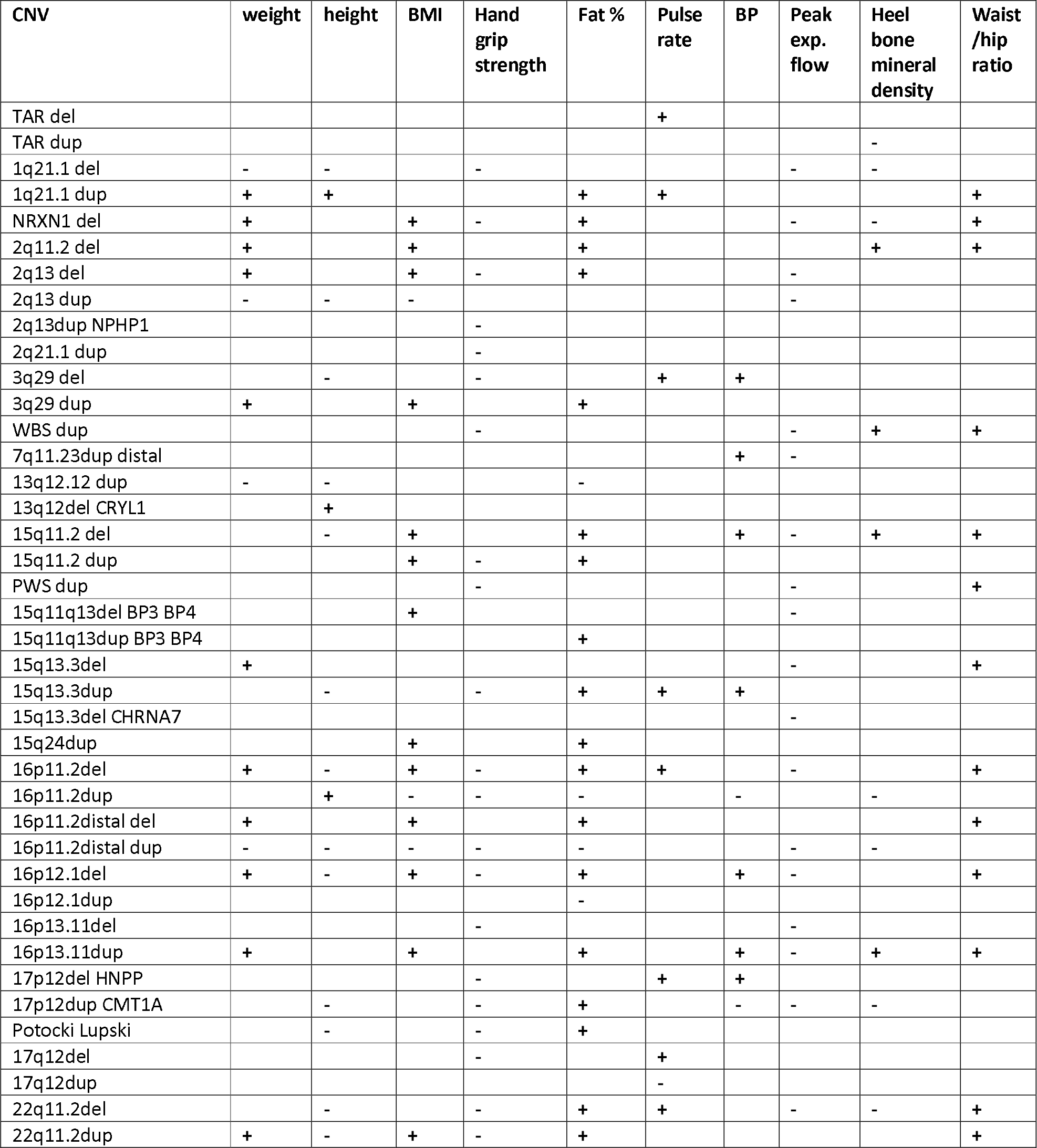
CNVs with significant associations at FDR=0.1 for physical traits. A “+” or “−” indicate the direction of the change. Fat % indicates a significant change on any of arm, leg or trunk fat % measurements.

| CNV | weight | height | BMI | Hand grip strength | Fat % | Pulse rate | BP | Peak exp. flow | Heel bone mineral density | Waist /hip ratio |
| --- | --- | --- | --- | --- | --- | --- | --- | --- | --- | --- |
| TAR del |  |  |  |  |  | + |  |  |  |  |
| TAR dup |  |  |  |  |  |  |  |  | - |  |
| 1q21.1 del | - | - |  | - |  |  |  | - | - |  |
| 1q21.1 dup | + | + |  |  | + | + |  |  |  | + |
| NRXN1 del | + |  | + | - | + |  |  | - | - | + |
| 2q11.2 del | + |  | + |  | + |  |  |  | + | + |
| 2q13 del | + |  | + | - | + |  |  | - |  |  |
| 2q13 dup | - | - | - |  |  |  |  | - |  |  |
| 2q13dup NPHP1 |  |  |  | - |  |  |  |  |  |  |
| 2q21.1 dup |  |  |  | - |  |  |  |  |  |  |
| 3q29 del |  | - |  | - |  | + | + |  |  |  |
| 3q29 dup | + |  | + |  | + |  |  |  |  |  |
| WBS dup |  |  |  | - |  |  |  | - | + | + |
| 7q11.23dup distal |  |  |  |  |  |  | + | - |  |  |
| 13q12.12 dup | - | - |  |  | - |  |  |  |  |  |
| 13q12del CRYL1 |  | + |  |  |  |  |  |  |  |  |
| 15q11.2 del |  | - | + |  | + |  | + | - | + | + |
| 15q11.2 dup |  |  | + | - | + |  |  |  |  |  |
| PWS dup |  |  |  | - |  |  |  | - |  | + |
| 15q11q13del BP3 BP4 |  |  | + |  |  |  |  | - |  |  |
| 15q11q13dup BP3 BP4 |  |  |  |  | + |  |  |  |  |  |
| 15q13.3del | + |  |  |  |  |  |  | - |  | + |
| 15q13.3dup |  | - |  | - | + | + | + |  |  |  |
| 15q13.3del CHRNA7 |  |  |  |  |  |  |  | - |  |  |
| 15q24dup |  |  | + |  | + |  |  |  |  |  |
| 16p11.2del | + | - | + | - | + | + |  | - |  | + |
| 16p11.2dup |  | + | - | - | - |  | - |  | - |  |
| 16p11.2distal del | + |  | + |  | + |  |  |  |  | + |
| 16p11.2distal dup | - | - | - | - | - |  |  | - | - |  |
| 16p12.1del | + | - | + | - | + |  | + | - |  | + |
| 16p12.1dup |  |  |  |  | - |  |  |  |  |  |
| 16p13.11del |  |  |  | - |  |  |  | - |  |  |
| 16p13.11dup | + |  | + |  | + |  | + | - | + | + |
| 17p12del HNPP |  |  |  | - |  | + | + |  |  |  |
| 17p12dup CMT1A |  | - |  | - | + |  | - | - | - |  |
| Potocki Lupski |  | - |  | - | + |  |  |  |  |  |
| 17q12del |  |  |  | - |  | + |  |  |  |  |
| 17q12dup |  |  |  |  |  | - |  |  |  |  |
| 22q11.2del |  | - |  | - | + | + |  | - | - | + |
| 22q11.2dup | + | - | + | - | + |  |  |  |  | + |

### Mirror phenotypes

A number of reciprocal deletions/duplications at the same locus show significant differences in opposite directions (as explained in the Discussion, no CNV is associated with increased physical strength or peak expiratory flow). We decided to use a simple definition for a CNV that produces a mirror phenotype: at least one measure should be changed in <u>opposite</u> directions, both <u>significantly</u> different from controls at FDR = 0.1. Using this definition, we find seven mirror image CNVs: 1q21.1, 2q13, 16p11.2 distal, 16p11.2, 16q12.1, 17p12 and 17q12 (Figure 2 a-g). Inspection of the directions of the effects of all CNVs (Supplementary Figure 1) suggests that more CNVs might produce true mirror phenotypes, if tested in larger samples. Our results confirm three of those reported by Macé et al, (13): 16p11.2 distal, 16p11.2 classic and 1q21.1, while 3q29 also suggests a mirror phenotype, but didn’t reach our criteria for inclusion, most likely due to the small number of observations. We note these two studies are not fully independent, as a third of the present sample was used in the earlier paper.

**Figure 2.**
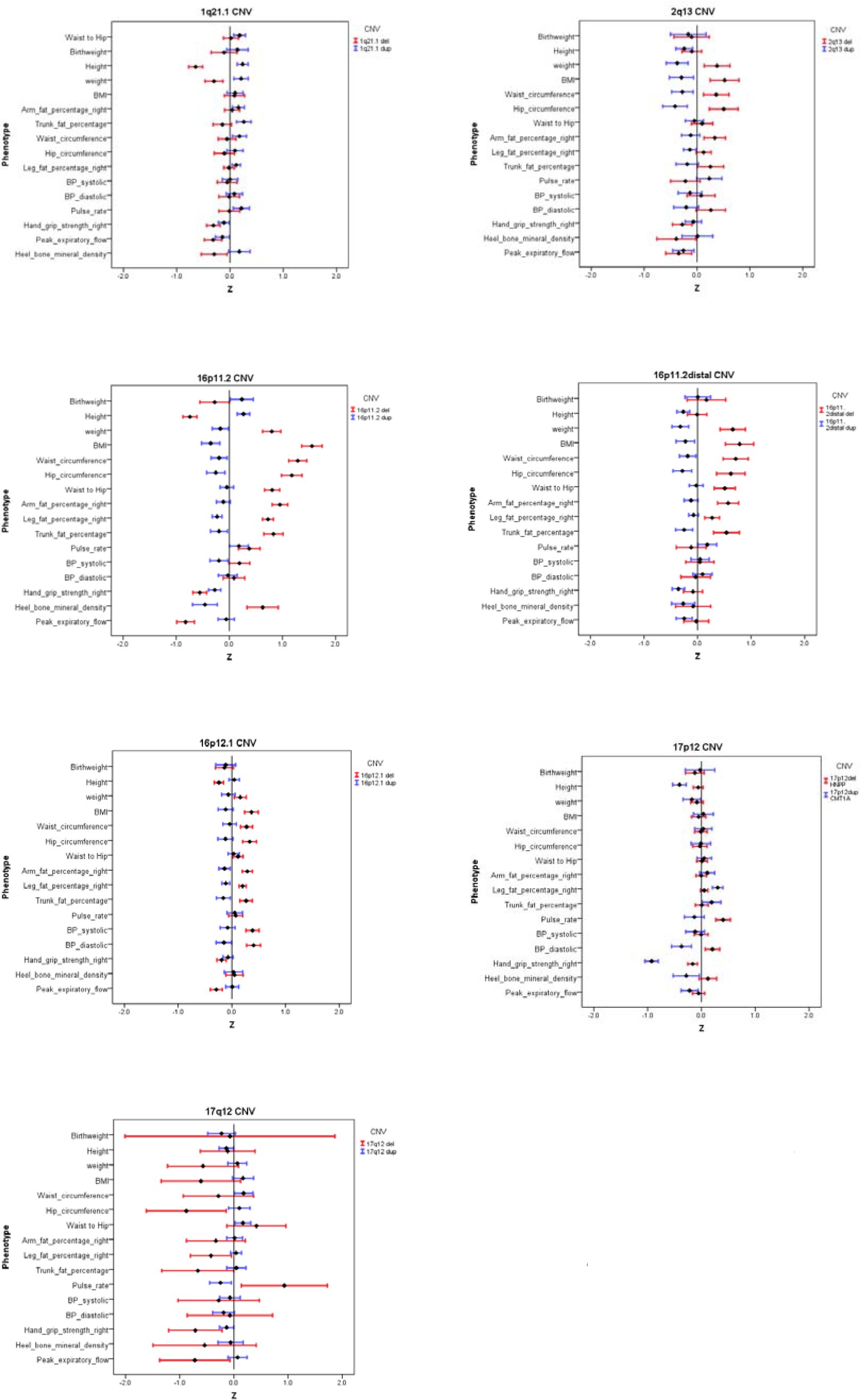
Effect sizes for the physical measurements for the seven CNVs with significant “mirror phenotypes”. a) 1q21.1, b) 2q13, c) 16p11.2, d) 16p11.2 distal, e) 16p12.1, f) 17p12, g) 17q12.

## DISCUSSION

### Novel associations

About one quarter of all possible CNV/phenotype associations were significant at FDR = 0.1, suggesting multiple effects on physical traits by many pathogenic CNVs. Some of these associations are already known from previous large case series (e.g. 16p11.2 deletion/duplication, 16p11.2 distal deletion), or from studies on individuals with syndromic short stature or obesity (Introduction), supporting the validity of this dataset and our methods. Many significant associations have not been reported before in systematically assessed cohorts, and certainly not as part of the same analysis, i.e. using identical methods to allow comparisons between CNVs. We show that the majority of these CNVs (41 of the 54) impact on at least one physical trait. For example, we find that deletions in *NRXN1*, a CNV so far known to increase risk for schizophrenia and autism spectrum disorders (20–22), are associated with a plethora of adverse changes, producing 11 significant results, such as increased weight, BMI, waist/hip ratio, fat percentage in arms, legs and trunk, and a faster pulse rate. *NRXN1* carriers also have reductions in muscular strength, peak expiratory flow and heel bone mineral density. Carrier status of 15q11.2 deletions and duplications have not been consistently associated with medical or anthropometric changes, although deletions increase risk for neurodevelopmental disorders (21, 23). We now show that deletion carriers have significant changes in over half of the measures, most notably reductions in height and birth weight, and increases in bone mineral density and waist/hip ratio. 15q11.2 duplication carriers have increased BMI, fat percentage and waist circumference and reduced muscular strength (Supplementary Table 1 and Figure 1). The magnitudes of the changes associated with 15q11.2 are small (e.g. deletion carriers are only 1.5cm shorter in height on average), but the very large sample size of the UK Biobank (~1600 deletion and ~1900 duplication carriers) allows small changes to be detected with high statistical confidence. Another notable finding is the association of 22q11.2 duplications with 10 measures. Although high rates of psychiatric disorders have been recorded for this CNV (24), increased weight has been reported only in single duplication carriers until now, and due to the extreme phenotypic variability of this condition, its true pathogenicity has been regarded as unclear (25). We report on a much larger sample (>260 carriers) and observe that central obesity (increased BMI and waist-to-hip ratio) is the leading feature in duplication carriers, making this CNV potentially highly pathogenic in relation to health-related outcomes (18,26).

### CNVs cause adverse effects of likely medical relevance

We note that most (although not all) of the observed effects are likely to be adverse for health. Thus, all 35 significant associations with physical strength and peak expiratory flow are in the direction of reduced performance. Both measures are associated with increased mortality (14,16). Most associations with pulse rate and blood pressure are in the direction of higher values, indicating a worse cardiovascular performance. Many CNVs cause increased weight/ fat percentage/ central obesity (increased waist/hip ratio), well-known factors increasing morbidity and mortality (15,18,26). There are notable exceptions: lower weight (and/or other obesity-related measures) are found in carriers of deletions at 1q21.1 and 16p11.2 distal, and duplications at 2q13, 13q12.12 and 16p11.2 classic. The potential protective effects of lower weight or fat percentage are however likely to be offset by other adverse consequences of these CNVs. Thirteen CNVs lead to shorter height, while only three are associated with increased height: 1q21.1 duplications and deletions at 13q12 (*CRYL1*) and 16p11.2 classic. BMI on its own is not sufficient to assess weight, height and obesity changes. For example, carriers of 1q21.1 deletions are on average shorter (−6.0cm) and lighter (−4.8kg), while carriers of the reciprocal 1q21.1 duplications are taller (+2.2cm) and heavier (+3.3kg).

Despite these substantial changes, both types of carriers have average BMI. A widely accepted measure of obesity, that has adverse effect on health outcomes, is the waist/hip ratio, which has been shown to better predict the risk for myocardial infarction (26). We find this significantly increased in 13 CNVs, including the 1q21.1 duplication, while no CNV caused a significantly reduced ratio, i.e. we do not see any protective effects for medical outcomes, even for CNVs that lead to reduced weight. Our findings that duplications at 15q11.2, 15q13.3, 16p13.3, 22q11.2 and deletions at *NRXN1* are among the CNVs with the highest rates of adverse changes on physical measurements, were unexpected and raise important issues regarding the management of such carriers.

### Are the effects primary or secondary?

Some of the changes in physical traits could be due to life-style differences among CNV carriers, or to concomitant medication given for diseases caused by the CNVs, rather than to a direct biological effect from gene dosage changes.

Current knowledge of CNV effects are largely restricted to childhood, while the UK Biobank population is composed of older adults, a difference that may result in lifestyle-related effects in this cohort. Reduced muscular strength, lower peak expiratory flow, faster pulse rate, increased weight and fat percentage can all be consequences of reduced exercise and life-style changes. Having a pathogenic CNV might make the person less likely to exercise, due to medical, cognitive or social problems. However, it appears that lifestyle is unlikely to account for all the changes we report. Thus, it cannot explain why at least seven CNV loci have opposite (mirror) phenotypes in deletion and duplication carriers (Figure 2) and why four CNVs lead to reduced weight (1q21.1 deletion, 2q13 duplication, 13q12.12 duplication and 16p11.2 distal duplication). The 1q21.1 duplication carriers present with increased height and weight, which together with the macrocephaly reported in children with this duplication (8) suggests that this is an overgrowth syndrome (27), although with a variable expressivity (8). Such observations indicate that many of the changes are a direct consequence of gene dosage changes, rather than secondary to life-style or social factors. In fact, each CNV has its own unique signature of physical traits, which become apparent on a heatmap image (Figure 3). Our work provides an unbiased comparison between these patterns among adults, as all Biobank participants were assessed with the same methods and blindly to their CNV status.

**Figure 3.**
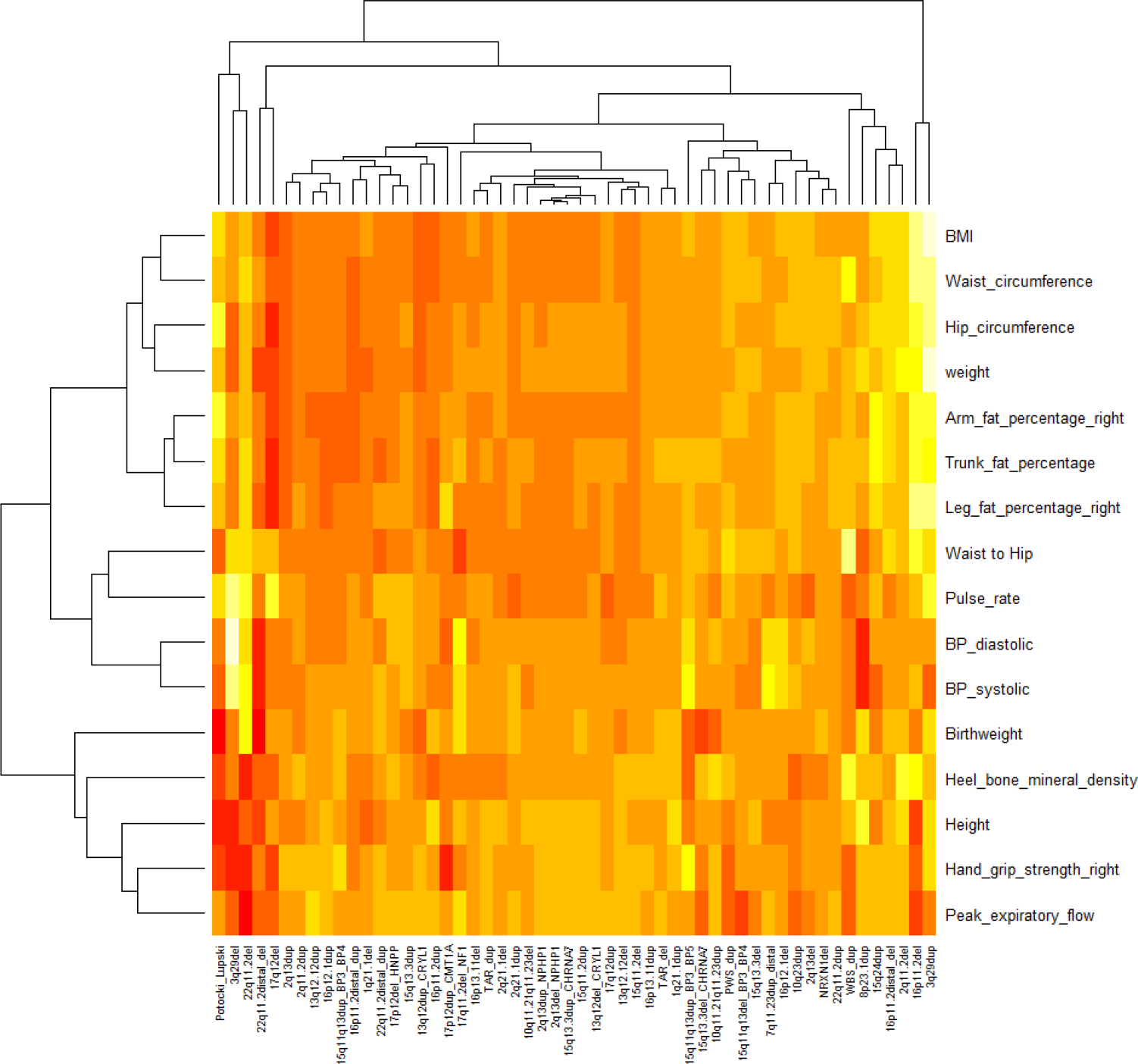
Clustering of the physical measurements for all 54 CNVs, with the normalised and corrected effect sizes (z-scores) of the 16 associations are clustered using the “heatmap” function of R. Paler colours indicate positive direction, while darker red show a negative direction of effect

### Conclusions

Our findings of adverse changes in basic physical characteristics in CNV carriers indicate that such individuals are at an increased risk for common medical disorders. Carriers of these CNVs could therefore benefit from general monitoring for cardiovascular risk factors and weight gain. It is tempting to envisage the targeting of genes or biochemical pathways in order to improve weight or fat distribution in humans. From this perspective, the CNVs with the most pronounced “mirror” phenotypes (Figure 2) are likely to contain the most promising candidate genes, as gene dosage changes lead to reciprocal changes in the measurements.

## MATERIALS AND METHODS

Approval for the study was obtained from the UK Biobank under project 14421: “Identifying the spectrum of biomedical traits in adults with pathogenic copy number variants (CNVs)”.

### CNV calling

We downloaded from the UK Biobank the raw intensity (CEL) files from Affymetrix BiLEVE (N~50,000) and Axiom (N~450,000) arrays and processed them with Affymetrix Power Tools (www.affymetrix.com/estore/partners_programs/programs/developer/tools/powertools.affx) and PennCNV (28). We followed the same CNV calling pipeline that we described previously (29). We then called a list of 92 CNVs proposed to be pathogenic, including their reciprocal deletions/duplications (23,30). After excluding CNVs with fewer than five observations, and three small CNV loci that produced false-positive calls (two of them telomeric), we took forward 54 CNVs for analysis of associations. Supplementary Table 2 lists the CNVs and the reasons for exclusion or retention in analysis.

### Physical measurements

We obtained data from the UK Biobank on tests, which were performed at the assessment centres and described as “physical measurements” (biobank.ctsu.ox.ac.uk/crystal/docs/Bodycomposition.pdf). These include anthropometric measures (height, weight, BMI, hip and waist circumference), body fat content, hand grip strength, spirometry, ultrasound heel bone densitometry and self-reported birthweight. Body fat content is estimated from the bioimpedance measures performed with a Tanita BC418MA body composition analyser. We also included pulse rate and blood pressure, as recorded at the assessment centres. In order to maximise statistical power and simplify the presentation of the results, we only used tests collected on >50% of individuals and excluded variables correlated at >0.9, such as measures collected on left and right arm or leg. We therefore only analysed measures performed on the right arm or right leg and averaged the two pulse rate measures that were performed at the same initial visit. Following research that highlights the importance of the waist to hip ratio (26), we added this measure too, resulting in a set of 16 variables (Table 2).

**Table 2.** List of physical measures from the UK Biobank data included in the analysis.

| **Measure** | **Biobank**<br>**Field ID** | **Procedure** | **Unit** | **Number of**<br>**people tested** |
| --- | --- | --- | --- | --- |
| Birthweight | 20022 | Self-report | kg | 225,138 |
| weight | 21002 | scales | kg | 395,584 |
| Height | 50 | Standing height | cm | 395,854 |
| BMI | 21001 | Weight/height<sup>2</sup> | kg/m<sup>2</sup> | 395,441 |
| Hand grip strength right | 47 | Grip strength device | kg | 395,129 |
| Waist circumference | 48 | Tape measure | cm | 396,037 |
| Hip circumference | 49 | Tape measure | cm | 395,992 |
| Waist to hip ratio | 48/49 | Waist/hip | ratio | 395,954 |
| Leg fat percentage right | 23111 | bioimpedance | % | 389,777 |
| Arm fat percentage right | 23119 | bioimpedance | % | 389,722 |
| Trunk fat percentage | 23127 | bioimpedance | % | 389,569 |
| Pulse rate | 102 | Automated reading | bpm | 373,789 |
| BP systolic | 4080 | Automated reading | mmHg | 370,501 |
| BP diastolic | 4079 | Automated reading | mmHg | 370,511 |
| Peak expiratory flow | 3064 | Spirometry | litres/min | 361,464 |
| Heel bone mineral density | 3148 | Ultrasound bone<br>densitometry | g/cm<sup>2</sup> | 228,405 |

### Statistical analysis

We filtered out poorly performing arrays using the following cut-off criteria: genotyping call rate < 0.96, > 30 CNVs per person, a waviness factor of < −0.03 & > 0.03 & LRR standard deviation of > 0.35. We excluded people who self-report to be other than white British or Irish and first-degree relatives (using the kinship coefficients data). This left 396,725 people for analysis. The tested variables followed normal distributions, therefore we did not further transform the data prior to analysis, other than to normalise all measures into z-scores, for a uniform presentation. We used linear regression analysis (glm) in R (version 3.3.2) to test the effect of the CNV carrier status on each measure (in z-score differences). For co-variates we used sex, age, array type (Axiom/BiLEVE), Townsend deprivation index (as a measure of the socioeconomic status) and the first 15 principal components from the genetic analysis, as provided by the UK Biobank. We also provide the changes in non-normalised (original) units, to give more real-world view of the effect of CNV carrier status (e.g. kg, beats per minute, mmHg). We did not control for education or occupation, as we have shown that these can be consequences of CNV carrier status (29). A Bonferroni correction for 864 test gives a level of significance of p < 5.8×10^−5^, which is conservative, given many of the measures are correlated (e.g. BMI and waist/hip ratio). We used instead the Benjamini-Hochberg FDR = 0.1 as our significance threshold (31), which corresponded to a nominal p-value of < 0.025 for the current study (Supplementary Table 1).

## Acknowledgements

This research has been conducted using the UK Biobank Resource under Application Number 14421.

